# Both rare and common genetic variants contribute to autism in the Faroe Islands

**DOI:** 10.1101/363853

**Authors:** Claire S Leblond, Freddy Cliquet, Coralie Carton, Guillaume Huguet, Alexandre Mathieu, Thomas Kergrohen, Julien Buratti, Nathalie Lemière, Laurence Cuisset, Thierry Bienvenu, Anne Boland, Jean-François Deleuze, consortium GenMed, Tormodur Stora, Rannva Biskupstoe, Jónrit Halling, Guðrið Andorsdóttir, Eva Billstedt, Christopher Gillberg, Thomas Bourgeron

## Abstract

The number of genes associated with autism is increasing, but few studies have been performed on epidemiological cohorts and in isolated populations. Here, we investigated 357 individuals from the Faroe Islands including 36 individuals with autism, 136 of their relatives and 185 non-autism controls. Data from SNP array and whole exome sequencing revealed that individuals with autism compared to controls had a higher burden of copy-number variants (*p* < 0.05), higher inbreeding status (*p* < 0.005) and higher load of homozygous deleterious variants (*p* < 0.01). Our analysis supports the role of several genes/loci associated with autism (e.g. *NRXN1, ADNP*, 22q11 deletion) and identified new truncating (*e.g. GRIK2, ROBO1, NINL* and *IMMP2L*) or recessive deleterious variants (*e.g. KIRELL3* and *CNTNAP2*) affecting autism-risk genes. It also revealed three genes involved in synaptic plasticity, *RIMS4, KALRN* and PLA2G4A, carrying *de novo* deleterious variants in individuals with autism without intellectual disability. In summary, our analysis provides a better understanding of the genetic architecture of autism in isolated populations by highlighting the role of both common and rare gene variants and pointing at new autism-risk genes. It also indicates that more knowledge about how multiple genetic hits affect neuronal function will be necessary to fully understand the genetic architecture of autism.

## Introduction

Autism spectrum conditions (ASCs; henceforth ‘autism’) are diagnosed in 1-2% of the population worldwide and are characterized by atypical social communication and the presence of restricted interests, stereotyped and repetitive behaviors. Individuals with autism can also suffer from psychiatric and medical conditions including intellectual disability (ID), epilepsy, motor control difficulties, attention-deficit hyperactivity disorder (ADHD), tics, anxiety, sleep disorders, depression or gastrointestinal problems (1). The genetic susceptibility to autism can vary from one individual to another. In some cases, a single *de novo* variant can be detected. On the contrary, in some cases, the genetic architecture is more complex and involved thousands of common genetic variants, each one with low impact but collectively increasing the susceptibility to autism (2). Most of our knowledge on the genetics of autism comes from studies on unrelated individuals with autism who do not share a recent common ancestor. Several studies investigated families with autism from countries where consanguinity is high (3,4), but the genetic architecture of autism in isolated populations remains largely unknown.

The Faroe Islands is an archipelago located in the North Atlantic Ocean, halfway between Norway, Iceland and Scotland (Fig 1A). The population (approximately 49,000 inhabitants) was founded in the 9^th^ century by a small number of emigrants from Norway. The population remained at a small size for centuries until they experienced a rapid expansion in the 1800s. Previous genetic studies indicated that individuals from Scotland, Norway, Sweden, Ireland, Iceland and British Isles have significantly contributed to the current gene pool of the Faroese population (5,6).

**Fig 1.**
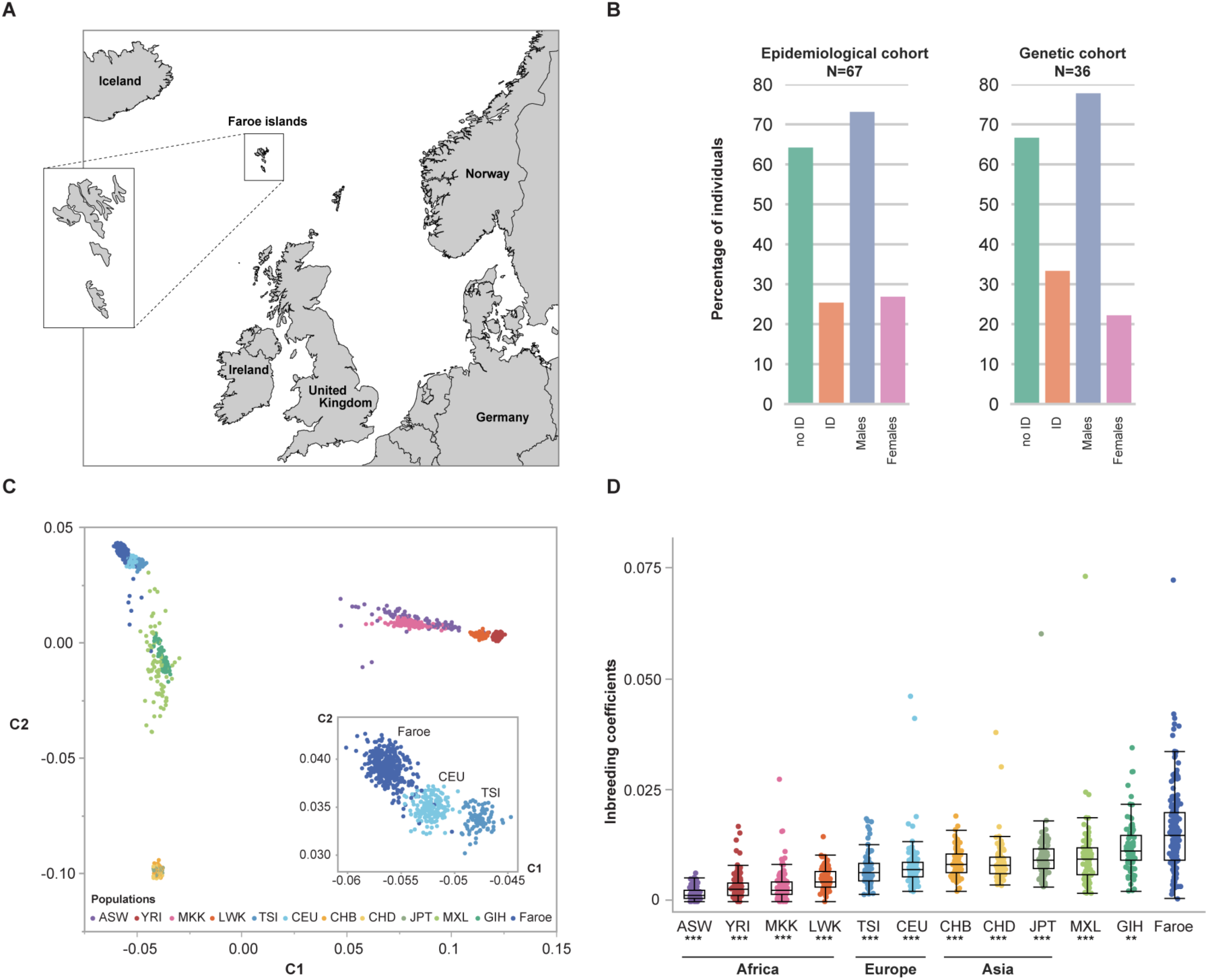
Genetic background of the Faroese population. A. Geographic localization of the Faroe Islands (R software and “maps” and “mapdata” packages were used to draw the map). B. Demographic comparison of the epidemiological and genetic cohorts of the Faroe Islands. The epidemiological and the genetic cohort are composed of 67 and 36 individuals with autism, respectively. C. Multidimensional scaling plots (MDS) of genome-wide identity by state (IBS) pairwise distances between human populations. Each dot represents an individual and the distance between two dots corresponds to genetic distance based on genome-wide pairwise IBS calculations. D. Degree of inbreeding across HapMap 3 and Faroese populations. The inbreeding coefficients of the Faroe non-autism control individuals (n=176) were compared to the HapMap3 populations. Faroe individuals displayed a higher degree of inbreeding compared with others human populations (Median F: F_FAROES_ = 0.015; F_ASW_ = 0.0014; F_CEU_ = 0.0071; F_CHD_ = 0.0081; F_CHB_ = 0.0083; F_GIH_ = 0.011; F_JPT_ = 0.0093; F_LWK_ = 0.0045; F_MXL_ = 0.0095; F_MKK_ = 0.0026; F_TSI_ = 0.0066; F_YRI_ = 0.0028; Paired samples Wilcoxon test: W_ASW_ = 12.56, *p*_ASW_ < 0.0001; W_CEU_ = 12.27, *p*_CEU_ < 0.0001; W_CHD_ = 7.14, *p*_CHD_ < 0.0001; W_CHB_ = 7.11, *p*_CHB_ < 0.0001; W_GIH_ = −3.06, *p*_GIH_ < 0.002;W_JPT_ = −5.83, *p*_JPT_ < 0.0001; W_LWK_ = −10.86, *p*_LWK_ < 0.0001; W_MXL_ = −5.66, *p*_MXL_ < 0.0001; W_MKK_ = −14.52, *p*_MKK_ < 0.0001; W_TSI_ = −8.64, *p*_TSI_ < 0.0001; W_YRI_ = −14.41, *p*_YRI_ < 0.0001; *p* are nominal p-values, ^*^*p* < 0.05, ^**^*p* < 0.01^, ***^*p* < 0.001). N, number; ID, intellectual disability; ASW, African ancestry in Southwest USA (n=83); CEU, Utah residents with Northern and Western European ancestry from the CEPH collection (n=165); CHB, Han Chinese in Beijing, China (n=84); CHD, Chinese in Metropolitan Denver, Colorado (n=85); GIH, Gujarati Indians in Houston, Texas (n=88); JPT, Japanese in Tokyo, Japan (n=86); LWK, Luhya in Webuye, Kenya (n=90); MXL, Mexican ancestry in Los Angeles (n=77), California; MKK, Maasai in Kinyawa, Kenya (n=171); TSI, Toscani in Italia (n=88); YRI, Yoruba in Ibadan, Nigeria (n=167).

We previously showed that the prevalence of autism in the Faroe Islands (0.94% of the population (7–9)) was similar to many other western countries. In this study, we ascertained the genetic profile of 357 individuals including an epidemiological cohort of 36 individuals with autism born between 1985 and 1994 (Fig 1 and S1 Fig), their relatives (n=136) and a group of 185 controls. We first investigated the known causes of autism and then identified new candidate genes. We also ascertained the impact of inbreeding and the load of deleterious homozygous variants on the risk of autism. Finally, both rare and common genetic variants were used to stratify individuals with autism and to compare their genetic and clinical profiles.

## Results

### The genetic diversity in the Faroe Islands

A total of 67 children and adolescents with autism were detected in a total population study of individuals aged 8-17 years living in the Faroe Islands and born between 1985 and 1994 (8,10). Thirty-six of the 67 individuals (54% of the total group) signed (or had parents who signed) informed consent forms and were included in the genetic study. Participants and non-participants in the genetic study were similar in terms of gender and cognitive abilities (Fig 1B). In addition, we collected DNA from 136 of their relatives and from 185 “non-autism controls. The genetic profile included a high-density Illumina SNP array interrogating > 4.3 millions of single nucleotide polymorphisms (SNPs) and a whole exome sequencing (WES) to discover new variants. Using identical-by-state (IBS) genomic distance (see materials and methods), we first compared the population structure of individuals from the Faroe Islands with worldwide populations (Fig 1C). All individuals were clustered in the Faroese population (with the exception of seven controls, but still with European genetic background). Using admixture, we showed that individuals from the Faroe Islands have their genome in majority constituted from “European component” (S2 Fig). As expected from the demographic history, individuals from the Faroese population displayed a higher degree of inbreeding compared with others world populations (Fig 1D).

### Contribution of *de novo* variants

We ascertained the burden of *de novo* variants since they are key players in the genetics of autism (11,12). The *de novo* variants were identified for 31 independent families including 28 individuals with autism and 45 siblings for whom DNA of both parents was available (see Clinical notes in S1 Appendix for the pedigrees). The combined analysis of genotyping and WES data revealed the presence of *de novo* chromosomal abnormalities and exonic copy*-*number variants (CNVs) in 3 out of 26 individuals with autism (11.5%) and 1 out of 43 siblings (2.3%). One female PN400129 had a trisomy of chromosome 21 and was diagnosed with autism, ID and Down syndrome (S3 Fig and S1 Table). One female PN400533 with atypical autism without ID carried a *de novo* 2.9 Mb deletion on chromosome 22q11 causing DiGeorge/velocardial syndrome. A male PN400115 with atypical autism without ID carried a *de novo* 425.5 kb deletion removing the six first exons of the *NRXN1α*.We also found a 91.4 kb deletion removing all exons of *ADNP* in a male with autism and ID (PN400125). The deletion was not found in the mother and was most likely *de novo*, but father’s DNA was not available and none of the SNPs within the deletion were informative to confirm the *de novo* status of the deletion. The *de novo* CNV observed in the sibling (PN400170) was a duplication of 782 kb affecting 5 genes (*CNPY1, DPP6, EN2, HTR5A, INSIG1* and *PAXIP1*).

Using the WES data, we detected the presence of *de novo* single nucleotide variants (SNVs) and small insertions/deletions (indels)(S2 Table). Overall, the rate of *de novo* exonic SNV/indels was similar to other studies (13) and was not different in individuals with autism (0.93) and their siblings (0.96). The variants were considered as probably deleterious when they were likely gene disruptive (LGD, for example stop gain or frame shift variant) or missense events with a combined annotation dependent depletion (CADD) score > 30 (MIS30) (14). There was also no significant increase in the rate of *de novo* deleterious variants in individuals with autism compared to their siblings and no significant enrichment in genes associated with autism (SFARI genes (15)) or expressed in the brain (Brain genes, see subjects and methods for gene selection). Nevertheless, several deleterious variants were identified in known genes for autism (*MECP2*) or compelling candidate genes (*RIMS4, KALRN, PLA2G4A)* (S4 Fig). Clinical details on the individuals with autism carrying those variants are available in the S1 Appendix.

### Contribution of rare CNVs and SNVs/indels variants

The overall burden of exonic CNVs was higher in individuals with autism compared to controls for both deletions (*p* = 0.02) or duplication (*p* = 0.006) (Fig 2A and S3 Table). The burden of deletions was also higher for autism-associated genes listed in the SFARI database (*p* = 0.01), for genes intolerant to loss-of-function variant (pLI > 0.9) (16) (*p* = 0.01) or genes expressed in the brain (*p* = 0.02). For duplications, only genes expressed in the brain were more frequently duplicated in individuals with autism compared to controls (*p* = 0.005). These differences however do not survive corrections for multiple tests and we had no significant difference between individuals with autism and their siblings. Among the SFARI genes affected by the CNVs, we identified a 58 kb maternal inherited deletion including the *IMMP2L*, a 2 Mb paternal inherited duplication on the pseudo-autosomal region 1 including *SHOX* and *ASMT*, and a 39 kb maternal inherited duplication of *TBL1XR1* (S3 Table).

**Fig 2.**
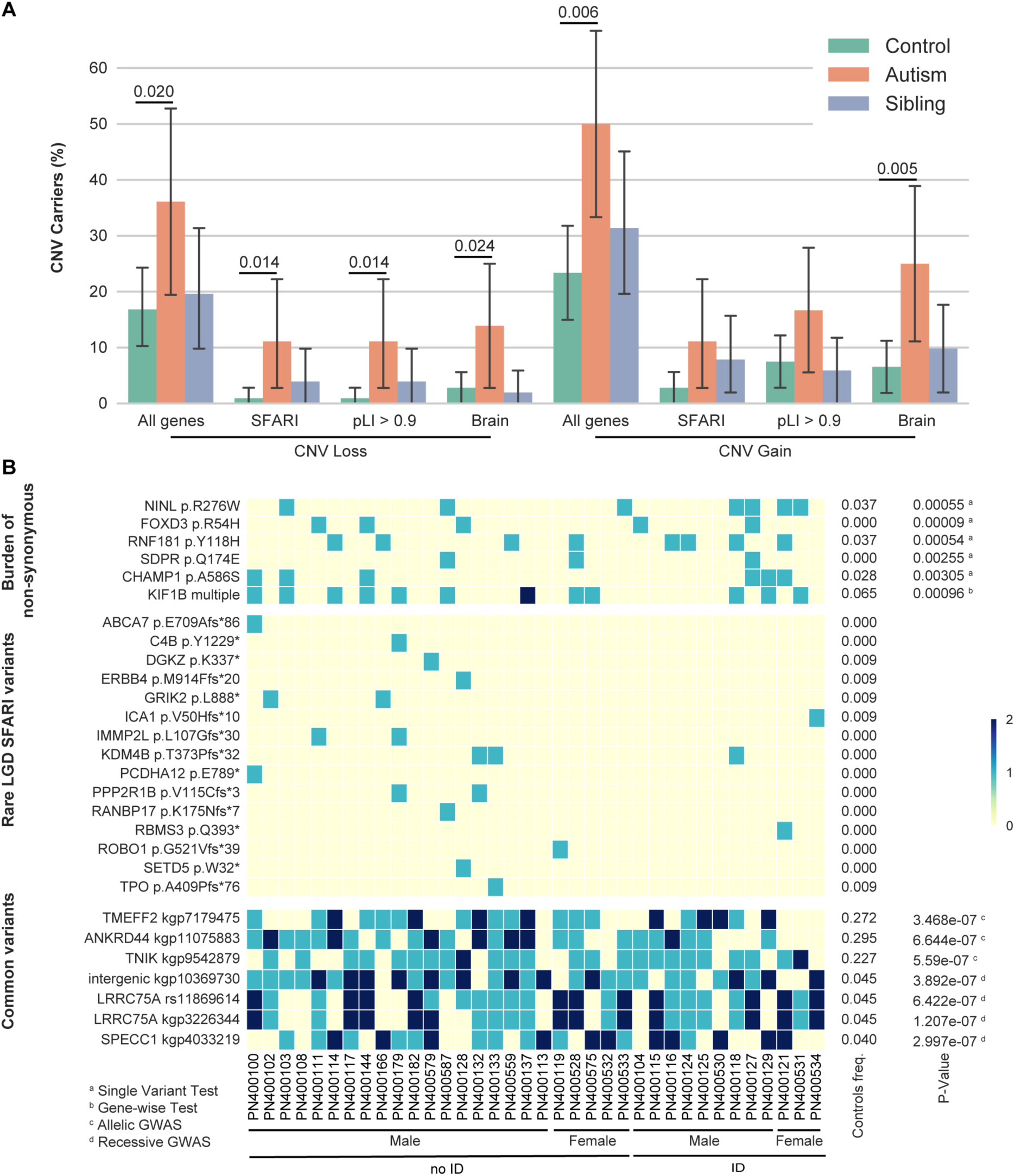
Rare and common variants in Faroese individuals with autism. A. Copy-number variant (CNV) analysis among gene-set lists within Faroe individuals. The number of exonic CNV carriers altering any genes or gene-set lists (SFARI genes, pLI > 0.9 genes and Brain genes, see Materials and Methods section) were compared between individuals with autism, siblings and controls (Fisher’s exact test: n_autism_ = 36, n_controls_ = 107, *p*_CNV_loss_All_genes_ = 0.02, OR_CNV_loss_All_genes_ = 2.79; *p*_CNV_loss_SFARI_ = 0.014, OR_CNV_loss_All_SFARI_ = 13.25; *p*_CNV_loss_pLI>0.9_ = 0.014, OR_CNV_loss_pLI>0.9_ = 13.25; *p*_CNV_loss_Brain_ = 0.024, OR_CNV_loss_All_Brain_ = 5.59; *p*_CNV_gain_All_genes_ = 0.0056, OR_CNV_gain_All_genes_ = 3.28; *p*_CNV_gain_SFARI_ = 0.067, OR_CNV_gain_All_SFARI_ = 4.33; *p*_CNV_gain_pLI>0.9_ = 0.11, OR_CNV_gain_pLI>0.9_ = 2.47; *p*_CNV_gain_Brain_ = 0.0049, OR_CNV_gain_All_Brain_ = 4.76; *p* are nominal p-values). B. Heatmap combining signals obtained from rare and common variant association tests and rare deleterious variants altering SFARI genes throughout Faroese individuals with autism. “Burden of non-synonymous” includes results from SKAT-O and CMC-EMMAX obtained from WES data (see Materials and Methods section and S5 fig; p < 10^^-3^). “Common variants” are the top hits of the genome wide association study (GWAS) for both allelic and recessive model obtained from genome-wide genotyping data (*p* < 10^^-6^). “Rare LGD SFARI variants” are rare likely gene disrupting (LGD) variants altering SFARI genes identified by (MAF < 1% in gnomAD). “The controls freq.” column indicates the proportion of non-ASD Faroese controls carrying the corresponding variant. P-Values are nominal. ID, intellectual disability.

Our analysis was restricted to CNVs affecting exons, but a large 357 kb duplication within intron 5 of the *NLGN1* gene and covering a long *NLGN1* antisense noncoding RNA was paternally inherited in a female (PN400102) with autism and no ID. There was no such rare intronic CNVs in the SFARI genes in siblings and controls.

We then run gene-wise association tests from rare SNV/indels (MAF < 5%). None of associations were gene wide significant, but *KIF1B, FOXD3, RNF181* and *SDPR* were among the top genes (*p* < 0.001) detected by both the CMC-EMMAX and SKAT-O analyses (Fig 2B, S5 Fig and S4-S5 Tables). For *FOXD3, RNF181*, and *SDPR the association was mainly* driven by one missense variant, whereas for *KIF1B*, several variants contributed to the association. Single variant tests were performed (S6 Table) and among the hits with *p* < 0.009, two variants affect genes previously associated with neurodevelopmental disorders (NDD). A variant p.R276W in *NINL*, a gene previously associated with autism (17) was more frequent in individuals with autism (P400121 was homozygous) compared to controls (*p* = 0.0005). This Ninein-like protein is known to associate with motor complex and interact with the ciliopathy-associated proteins lebercilin, USH2A and CC2D2A. A *CHAMP1* variant (p.A586S) was more frequent in individuals with autism compared with controls (*p* < 0.003) and it was previously shown that *de novo* variants in *CHAMP1* cause ID with severe speech impairment (18).

Finally, we found truncating variants affecting several SFARI genes (*e.g. PRODH, ERBB4, GRIK2, ROBO1, RBMS3*, and *IMMP2L*; Fig 2B and S7 Table), but the number of individuals with autism carrying these variants was too low to detect a significant association.

### Contribution of recessive variants

Since inbreeding increases the risk for individuals to carry homozygous deleterious variants, we first compared the inbreeding status of the individuals with autism, their relatives and controls. Patients (*p* < 0.005) as well as their siblings (*p* < 0.01) had a higher inbreeding coefficient compared with controls (Fig 3A). More interestingly, we found that individuals with autism were carrying more deleterious homozygous variants (LGD, MIS30, gnomAD MAF < 1%) than controls (*p* < 0.0005; Fig 3B and S8 Table). Genes carrying deleterious homozygous variants in affected individuals were significantly enriched in the combined gene-set lists (SFARI + pLI > 0.9 + Brain genes) compared to controls (*p* < 0.05; Fig 3C).

**Fig 3.**
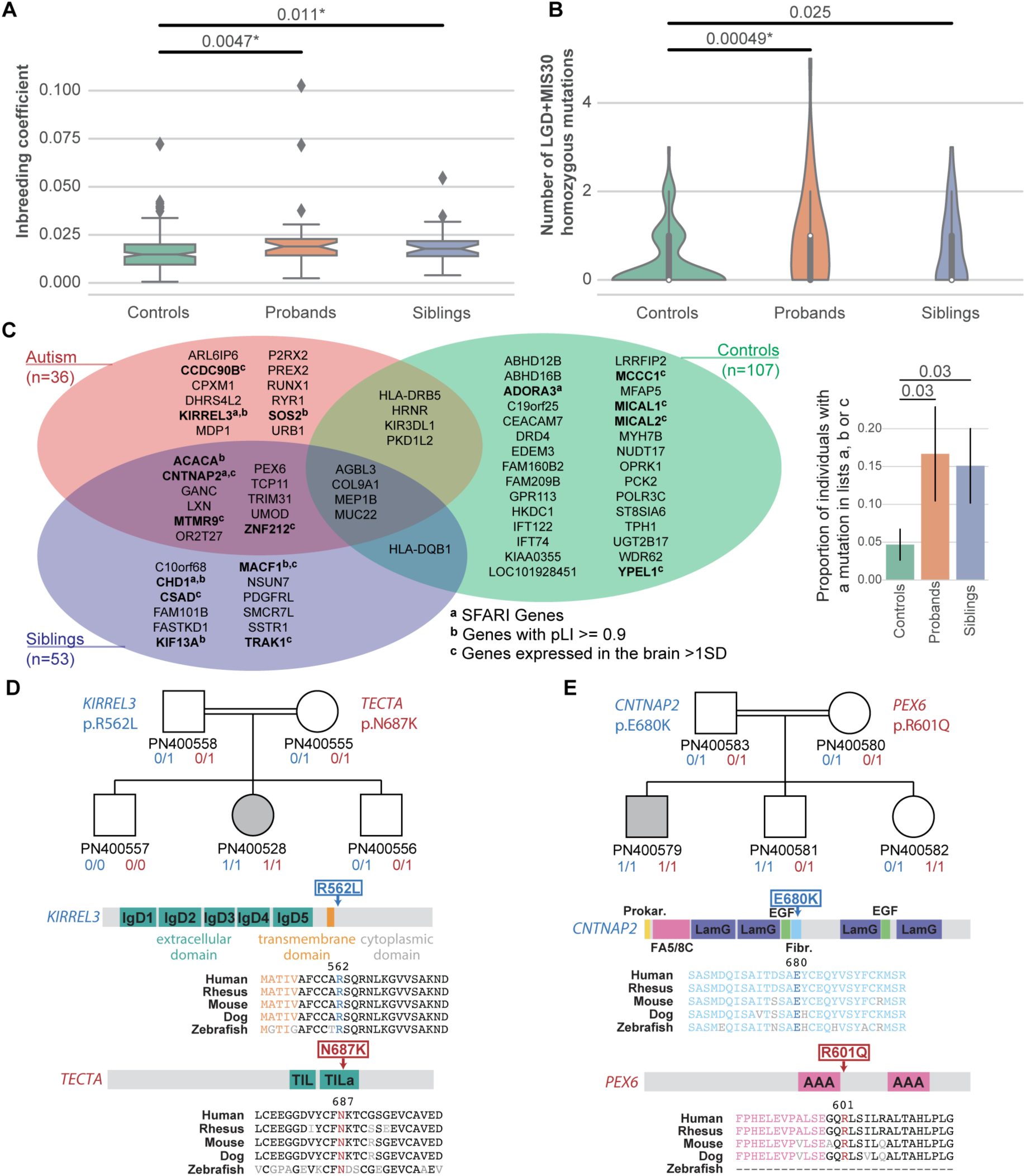
Genetic recessive mutations in Faroese individuals with autism. A. Distribution of the inbreeding coefficient in Faroese individuals (Mann Whitney U-test: n_autisms_ = 36, n_controls_ = 176, n_siblings_ = 30; U_controls.vs.autisms_ = 2,297, *p*_controls.vs.autisms_ = 0.0047; U_controls.vs.siblings_ = 1,953, *p*_controls.vs. siblings_ = 0.011; * indicates the one withstanding multiple testing). B. Number of rare LGD+MIS30 homozygous mutations carried per individual (Mann Whitney U-test: n_autisms_ = 36, n_controls_ = 107, n_siblings_ = 30; U_controls.vs.autisms_ = 1,321, *p*_controls.vs.autisms_ = 0.00049; U_controls.vs.siblings_ = 1,293, *p*_controls.vs. siblings_ = 0.025; * indicates the one withstanding multiple testing). C. Venn diagram of the genes carrying the variants from B. Genes names are in bold and annotated when they are part of our gene-set lists (SFARI genes, pLI > 0.9 genes and Brain genes, see subject and methods section). The plot on the right shows the proportion of individuals in each category carrying at least one mutated gene in our gene-sets lists (Fisher’s exact test: *p*_controls.vs.autisms_ = 0.03; *p*_controls.vs. siblings_ = 0.03). D. and E. are describing two specific families carrying multiple variants. “0” and “1” refer to wildtype or mutated allele, respectively. The localizations of the variants are indicated along the proteins and alignments throughout species showed the strong conservation of the altered amino acids. LGD, likely gene disruptive; MIS30, missense variants with CADD score ≥ 30; IgD, immunoglobulin domain; TIL, Trypsin Inhibitor-like; FA5/8C, Coagulation factor 5/8 type C domain; LamG, Laminin G domain; EGF, epidermal growth factor like domain; Fibr., Fibrinogen, alpha/beta/gamma chain, C-terminal globular domain; AAA: ATPases associated domains.

In one consanguineous family, we found a *KIRREL3* homozygous damaging missense variant (p.R562L) affecting a conserved residue in the cytoplasmic domain of this synaptic adhesion molecule (19) listed in SFARI and associated with NDD (20,21) (22) (Fig 3D). Interestingly, the female PN400528 with autism and a normal IQ was also homozygous for another deleterious variant (p.N687K) affecting *TECTA*, a SFARI gene associated with autism (21) and deafness (23).

In another family, the male PN400579 was homozygous for two variants affecting *CNTNAP2* and *PEX6* (Fig 3E). Recessive *CNTNAP2* variants are associated with Pitt-Hopkins like syndrome 1 and cortical dysplasia-focal epilepsy syndrome (MIM #610042). The *CNTNAP2* p.E680K variant affects a highly conserved amino acid within the fibrinogen domain of the protein. Recessive *PEX6* variants are associated with Heimler syndrome 2 a recessive peroxisome disorder characterized by sensorineural hearing loss, amelogenesis imperfecta and nail abnormalities, with or without visual defects (MIM #616617). The homozygous variant p.R601Q carried by the male with autism can be considered pathogenic since it was previously detected in several independent patients diagnosed with Heimler syndrome 2 (24). Details on the clinical profiles of the families are available in S1 Appendix.

### Contribution of common variants

In order to complete the genetic profile of all the individuals, we investigated the contribution of the common variants (MAF > 5%) using three complementary approaches: (i) genome wide association studies (GWAS) in cases and controls using three models (allelic, recessive and dominant), (ii) a burden/collapsing test that aggregates the variants located in a gene, (iii) a calculation of the autism polygenic risk score.

The results of the GWAS are presented in S6-S7 figs and a list of the top 45 loci that display p-values under 10^−5^ are shown in Fig 2B and S9 Table. Two of these loci were located within or nearby genes previously associated with NDD. *TNIK* on chromosome 3q26 (rs1492859; *p* = 5.59×10^−7^) encodes a key synaptic partner of *DISC1*, a gene associated with psychiatric disorders (25). *TMEFF2* on chromosome 2q32.3 (rs6737056; *p* = 3.47×10^−7^) is highly expressed in the brain and is known to modulate the AKT/ERK signaling pathways that are supposedly perturbed in autism (26). *TMEFF2* is also a target of *CHD8*, a chromatin remodeling gene associated with autism (27,28). Using the gene-based analysis implemented in MAGMA (29), only one gene *WHAMM* on chromosome 15q25.1 had a *p* < 10^−5^ (S10-S11 Tables). *WHAMM is expressed in the brain and acts as a regulator of membrane dynamics that* functions at the interface of the microtubule and actin cytoskeletons (30).

Finally, we ascertained the autism polygenic risk score (PRS-autism) for each individual (Fig 4). The PRS-autism was calculated using PRSice-2 from a previous GWAS using over 16,000 individuals with autism (31) who do not overlap with this sample. We found a significantly higher PRS-autism in individuals with autism compared to controls (*p* < 0.01; Fig 4A). Remarkably, in the autism group, the PRS-autism was significantly higher in individuals without ID compared to those with ID (*p* < 0.05, Fig 4B).

**Fig 4.**
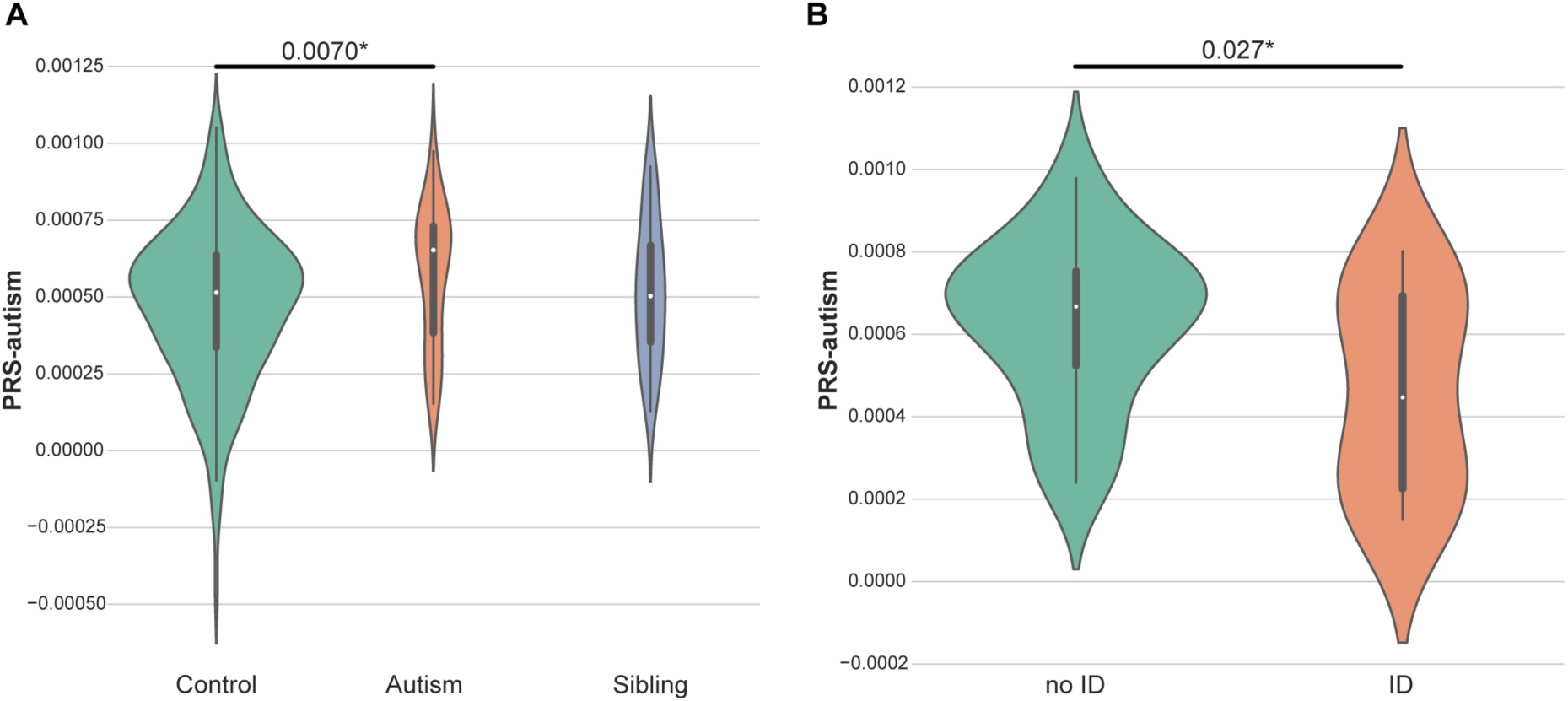
Distribution of the polygenic risk score for autism in Faroese individuals. A. Distribution of the polygenic risk score for autism (PRS-autism) of controls, autisms and siblings (Mann Whitney U-test: n_autisms_ = 36, n_controls_ = 107, n_siblings_ = 53; U_controls.vs.autisms_ = 2,344, *p*_controls.vs.autisms_ = 0.0070; * indicates the one withstanding multiple testing). B. Distribution of the PRS-autism for the cases without intellectual disability (ID) and the cases with ID (Mann Whitney U-test: n_autis ms-with-ID_ = 12, n_autisms-without-ID_ = 24; U_ID.vs.no-ID_ = 86, *p*_ID.vs.no-ID_ = 0.027; * indicates the one withstanding multiple testing). The PRS was calculated using PRSice-2 (see Materials and Methods section).

### Genetic stratification of autism in the Faroe Islands

In order to stratify individuals with autism, we used the number of rare deleterious variants in SFARI genes (including CNVs) and the PRS-autism estimated from the common variants (Fig 5). Hierarchical clustering found four clusters. The first one comprised seven individuals with high PRS-autism and high burden of deleterious variants in SFARI genes. In this cluster, none of the individuals had ID. The second “cluster” included only one individual who was diagnosed with autism and Down syndrome with a low PRS-autism and high burden of SFARI deleterious variants. In the third cluster, fourteen individuals had low PRS-autism and low burden of SFARI genes deleterious variants. In this cluster 50% of the individuals had ID. In the fourth cluster, fourteen individuals had higher PRS-autism, but low burden of SFARI deleterious variants. In this cluster, 29% of the individuals had ID and it includes the three individuals with autism with epilepsy from the cohort.

**Fig 5.**
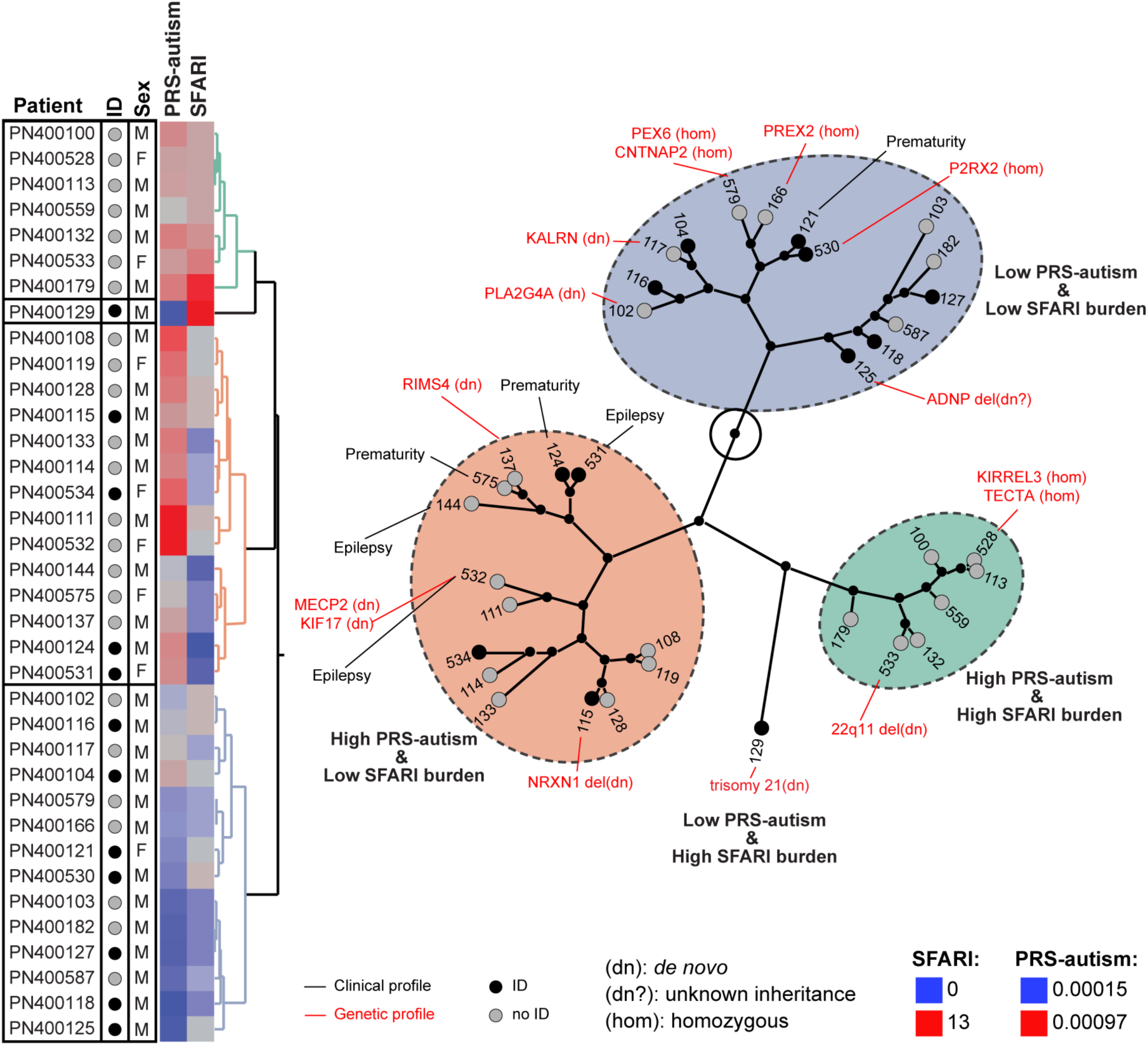
Stratification of autism in Faroese individuals. On the left, the stratification was built using hierarchical clustering on the number of genes carrying rare deleterious variants altering SFARI genes (MIS30, LGD or CNV) and on the polygenic risk score for autism (PRS-autism). The other columns were not used for the clustering. The genetic profile contains variants with a predicted impact on the condition of the individual with autism, the one in bold are most likely causatives. The clinical profile gives a subset of relevant information for each individual with autism. ID, intellectual disability; M, male; F, female; del, deletion; dup, duplication.

## Discussion

In this study, we investigated a group of individuals with autism that has two distinctive features. First, the group is representative of a general population cohort of all young people living in the Faroe Islands at one point in time, meaning that it was not biased for inclusion/exclusion criteria used for research studies. Secondly, the Faroese population has a more homogeneous genetic background compared to most other populations.

### Autism-risk genes in the Faroe Islands

We found a subset of individuals carrying strongly deleterious variants (some of which appeared *de novo*) affecting single gene or chromosomal regions. The chromosomal abnormalities included on case each of trisomy 21 and 22q11 deletion (causing Down and DiGeorge/velocardial syndromes, respectively). This was not surprising to find such known genetic disorders in an epidemiologic cohort since the prevalence of autism in individuals diagnosed with these syndromes is higher than in the general population (16-37% for Down syndrome (32), and 23-50% for DiGeorge/velocardial syndrome deletion (33,34)). We also revealed new variants in known autism-risk genes (*ADPNP, NRXN1, NINL, MECP2*) and points at new compelling candidate genes such as *KALRN, PLA2G4A* and *RIMS4*.

The *KALRN* gene encodes for a guanine nucleotide exchange factor (GEF) with strong homology to *TRIO,* a gene previously associated with autism (35). *KALRN* is expressed in neuronal tissue during embryonic development (36) and has been associated with schizophrenia risk through association analysis and postmortem analyses of individuals with autism’ cortical KALRN mRNA and protein levels (37),(38). It is also a binding partner of the Huntingtin and a regulator of structural and functional plasticity at dendritic spines. The *de novo* variant (p.N2024D; CADD=26.7) was never observed in the general population affects a key amino acid of the GEF domain conserved through evolution and present in *Drosophila melanogaster* and *Caenorhabditis elegans*. The male individual (PN400117) carrying this *de novo KALRN* variant has no ID.

*PLA2G4A* gene encodes the cytosolic phospholipase A2α that catalyzes the hydrolysis of membrane phospholipids to produce arachidonic acid. Mice lacking *Pla2g4a* display abnormalities in neuronal maturation (narrow synaptic cleft) (39) and long-term potentiation (LTP) (40). The *de novo* variant (p.R485C; CADD=35) has never been observed in the general population and is predicted as a deleterious variant falling in the catalytic domain of the protein. The female PN400102 carrying this *de novo variant has no ID.*

*RIMS4* codes for a presynaptic proteins that plays a key role for dendritic and axonal morphogenesis (41). *RIMS1* and *RIMS3* have already been associated with autism (42–44). The individual (PN400137) carrying the *de novo RIMS4* stop variant (p.Y204*; CADD=38) has a normal IQ (Performance IQ=108, Verbal IQ=116). The variant is predicted to truncate the last quarter of the protein and was never observed in the general population. Interestingly, RIM proteins interact with voltage-dependent Ca(2+) channels (VDCCs) and suppress their activity at the presynaptic active zone to regulate neurotransmitter release. Knockdown of gamma-RIMs (*RIMS3* and *RIMS4*) attenuated glutamate release to a lesser extent than that of alpha-RIMs (*RIMS1* and *RIMS2*). As a consequence, competition between alpha- and gamma-RIMs seems to be essential for modulating the release of glutamate at the synapse (45). We can therefore hypothesize that the *de novo RIMS4* truncating stop variant perturbs the fine-tuning of glutamatergic release at the synapse and contributes to autism.

### High inbreeding slightly increases the risk of autism, but no evidence for a founder effect for autism in the Faroe Islands

In genetic isolates, it is frequent to observe an increase frequency of diseases due to the presence of deleterious variants that were present in the genomes of the small group of migrants who settled the population. In the Faroe Islands, this “founder effect” was documented for several genetic diseases such as Bardet-Biedl syndrome (46), cystic fibrosis (47), 3-Methylcrotonyl-CoA carboxylase deficiency (48), glycogen storage disease type IIIA (49) and retinitis pigmentosa (50). We confirmed that inbreeding in the Faroe Island is higher than expected compared with other populations. The median of inbreeding is F = 0.015 ± 0.001 in the control sample and is similar to the one reported by Binzer et al. (2014) in their study on multiple sclerosis in the Faroe Islands (F = 0.018) (51). This level of inbreeding corresponds approximately to children from parents with a second-cousin relationship (F = 0.016). We also observed that individuals with autism from the Faroe Islands have significantly higher level of inbreeding and burden of recessive deleterious variants compared to their geographically matched controls. The homozygous deleterious variants carried by individuals with autism were enriched in genes included in our list of genes of interest (*e. g.* high intolerance for loss of function variants, expressed in the brain and previously associated with autism). However, one should note that the increased probability for having autism due to inbreeding in the Faroe Islands is relatively small (F_autism_ = 0.0189; F_controls_ = 0.0148; OR = 1.28; *p* = 0.0047).

In contrast to other genetic conditions, we could not detect a founder effect for autism in the Faroe Islands. Moreover, the loci identified in our study do not overlap with those detected in a previous genetic microsatellite association study in the Faroese population pointing at regions on 2q, 3p, 6q, 15q, 16p, and 18q (52). We also found no overlap between the variants identified in our study and those found in Faroese individuals with autism diagnosed with panic (53) or bipolar (54) disorders. This absence of a founder effect is also in agreement with the epidemiological observation that the prevalence of autism in the Faroese population is not higher compared to more outbred populations.

### Perspectives

Our study confirms that both rare and common genetic variants contribute to the susceptibility to autism. Indeed, although, we identified previously known genetic causes for autism and pointed at new compelling candidate genes, we also showed a contribution of the common variants illustrated by the higher level of PRS-autism in individuals with autism (especially those with no ID) compared to controls. To date, in the literature, very few genes are identified in individuals diagnosed with autism, but with intact general intelligence. Based on the genes previously reported (*NLGN3, NLGN4X*, duplication of *SHANK3*) (55,56) and the genes found in this study (*RIMS4, KALRN, PLA2G4A*), it seems that the proteins involved in autism without ID converge to different part of the post- and pre-synapse rather than to pathways such as gene regulation and chromatin remodeling, but this has to be confirmed on larger cohorts. Indeed, the main limitation of our study is the small number of individuals with autism. Several LGD variants affecting autism-risk genes such as *GRIK2* (57) or *ASMT* (58) were found exclusively or more frequently in individual with autism compared to controls, but we lack a replication samples to confirm the contribution of these variants in the susceptibility to autism in the Faroe Islands.

In summary, this study improves our knowledge on the genetic architecture of autism in epidemiological cohorts and in genetic isolates by showing that the contribution of both rare and common gene variants to autism can be detected in small, but genetically homogeneous populations. It also provides new compelling candidate genes and reveals that high inbreeding and high load of homozygous deleterious variants can be a risk factor for autism. Such combined analysis investigating both rare and common gene variants might represent a useful framework to investigate, from groups to individuals, the complex genetic architecture of autism.

## Materials and Methods

### Ethics statement

This study was approved by the IRB of the “Institut Pasteur” of Paris (IRB00006966 Institut Pasteur, approval 2010-003).

### Patients

All individuals with autism in this study were recruited from an epidemiological cohort through a two-phase screening and diagnostic process targeting all children born in the 10-year period from 1985 through 1994 and living in the Faroe Islands in 2002 (7-16 years, n=7,689 children) and 2009 (15-24 years, n= 7,128 children) (8),(10). The total number of children diagnosed with autism was 67 which corresponds to an autism prevalence of 0.94%. Among the individuals with autism, 23% were diagnosed with childhood autism, 56% with Asperger syndrome and 21% with atypical autism. There were 49 males (73.1%) and 18 females (26.9%). DNA was available for 36 individuals with autism including 11 diagnosed with childhood autism (31%), 17 with Asperger syndrome (47%), and 8 with atypical autism (22%). There were 28 males (78%) and 8 females (22%). The non-autism controls were recruited by issuing an invitation with information on the study to all pupils at the high school level in winter 2008-2009. The schools invited are in Eysturoy, Suduroy and Torshavn. The age of the invited was from 14-24 years. For those under 18 years a letter was sent to the parents that could sign the consent for their children.

### Screening and diagnosis

In 2002, screening included the use of the Autism Spectrum Screening Questionnaire (ASSQ) (59). Screen-positive children were thoroughly examined via Diagnostic Interview for Social and Communication Disorder (DISCO-10 in 2002 and DISCO-11 in 2009) (60) of one or both parents and the Wechsler Intelligence Scale for Children – 3^r^ edition (WISC) or Wechsler Adult Intelligence Scale – Revised (WAIS). Whenever overall and verbal abilities allowed it possible, children were also interviewed in an unstructured/semistructured manner about interests and skills patterns, peer relations, family relationships and about formal general information knowledge. The following diagnostic criteria used when making clinical diagnoses were (a) ICD-10 criteria for childhood autism/autistic disorder; (b) Gillberg criteria for Asperger syndrome; (c) ICD-10 criteria for atypical autism with the added requirement that a case thus diagnosed could not meet full criteria for childhood autism or Asperger syndrome; and (d) ICD-10 criteria for disintegrative disorder.

The majority of children in the atypical autism and Asperger syndrome groups had been tested with the WISC-R. Those with childhood autism had usually been tested on other tests. In those intellectually low-functioning individuals for whom no test was available, IQ was estimated on the basis of the Vineland developmental portion that is part of the DISCO-interview.

### Genotyping

The cohort available for the genotyping is shown in Fig 1 and S1 Fig. It includes 36 individuals with autism, 208 controls, 132 close relatives of the individuals with autism (61 siblings and 68 parents) and 10 close relatives from the controls. DNA was extracted from blood leukocytes. The genotyping was performed at the “Centre National de Recherche en Génomique Humaine (CNRGH)” using the Infinium IlluminaOmni5-4 BeadChip (> 4.3 millions of markers) from Illumina. Sample quality controls such as Sex check (based on the X chromosome homozygosity rate), Mendel errors (transmission errors within full trios) and Identity By State (IBS, see section below) were performed using PLINK 1.90 (61).

### Population genetic structure

Genome-wide pairwise IBS calculations and Multidimensional scaling (mds) analysis on genome-wide IBS pairwise distances matrix was calculated using PLINK 1.90. IBS values have been calculated for 376 individuals from Faroe Islands and 1,184 individuals from HapMap phase 3 project with the following calculation: 1- (0.5 * IBS1 + IBS2)/N; N is the number of tested markers; IBS1 and IBS2 are the number of markers for which one pair of individuals share either 1 or 2 identical allele(s), respectively. Out of the 376 individuals, 32 individuals were removed from further analyses, including 7 ancestry outliers (all controls), 9 siblings of controls, one swap and 15 control individuals involved in pairs with IBS score superior to 0.9.

For the estimation of the inbreeding status, SNPs with genotyping call rate < 95%, minor allele frequency < 0.05, strong linkage disequilibrium r > 0.5 or failing Hardy Weinberg equilibrium test (*p* < 10^−6^) were filtered out of the Faroe SNP genotyping dataset. All homozygosity analyses were performed with Plink 1.09 on autosomes including identification of Runs Of Homozygosity (ROH) and Inbreeding coefficients calculation. For ROH detection, a threshold of 50 consecutive homozygous SNPs with a minimum density of 1 SNP / 5,000 kb and no minimum length 50 SNPs was used following Howrigan et al.’s guidelines (62). No heterozygous markers were allowed in the 50 SNPs-window. In this analysis, the maximum gap between two consecutive SNPs within a run was set to 5,000 kb. Inbreeding coefficients were calculated by estimating the proportion of the autosomal genome that is in ROH. This method was proposed by McQuillan and al (2008)(63) and has been showed to be the most reliable, especially with small sample size(64). Faroe inbreeding coefficients were compared to inbreeding coefficient of HapMap phase 3 project populations.

### Genome-wide association study (GWAS)

Prior association analyses, SNPs with genotyping call rate < 90%, minor allele frequency < 0.05 or failing Hardy Weinberg equilibrium test (*p* < 10^−6^) were filtered out of the Faroe SNP genotyping dataset. The global genome wide genotyping call rate of all the individuals was superior to 90%. A total of 1,690,491 variants and 212 independent individuals (including 36 cases and 176 controls) passed filters and QC. Allelic, recessive and dominant GWAS were performed with Plink 1.09 using Chi-squared statistics. Manhattan and Quantile-Quantile (Q-Q) plots were generated using R. Gene and gene-set (including SFARI, pLI > 0.9 and Brain gene lists) analyses were performed with MAGMA v1.06 (29) using principal components regression and linear regression model, respectively.

### Polygenic risk score (PRS) for autism

The computation of the PRS was performed with the tool PRSice2 (65) on the Faroes SNP array data using as a reference the PGC GWAS summary statistics (31). SNPs were not imputed since we used high density arrays (over 4 millions SNPs). For our dataset, PRSice2 with default parameters defined a p-value threshold of 0.197 which gives us a R^2^ (squared correlation coefficient) of 0.04.

### Whole-Exome Sequencing (WES)

Blood leukocytes DNA from 286 individuals was enriched for exonic sequences through hybridization SureSelect Human All Exon V5 (Agilent) by the CNRGH. For 67 individuals for whom the available quantity of DNA was low, they used a low-input protocol using only 200 ng of DNA compared to 3 µg for the normal protocol. The captured DNA was sequenced using a HiSeq 2000 instrument (Illumina). Coverage/depth statistics have been accessed as quality control criteria. We required that more than 90% of each exome have 10X coverage and more than 80% have 20X coverage. Short read sequences were then aligned to hg19 with BWA v0.7.8, duplicate reads were removed with PicardTools MarkDuplicates. Reads with a global quality under 30 and a mapping quality under 20 were excluded from the analysis. Variants were predicted using FreeBayes (66) and GATK (67) with a minimum of 10 reads covering the position. VEP (using RefSeq and Ensembl 91) was used to annotate the variants. We used the GEMINI (68) framework that automatically integrates the VCF file into a database for exploring genetic variant for disease and population genetics. Genetic variants were analyzed using GRAVITY, a Cytoscape (69) plugin designed in the lab specifically for interpreting WES results using Protein-Protein Interaction networks (http://gravity.pasteur.fr/). Gravity uses a user-friendly interface and makes easier prioritizing variants according to damage prediction, mode of inheritance, gene categories and variant frequency in databases. It allows filtering variants with many parameters, such as quality parameters (DQ, MQ, GQ), allelic fraction, frequency of the variant in the cohort or in databases, damage prediction scores (CADD, SIFT, Polyphen2) and many more. Since WES does not detect the *FMR1* amplification, 33 individuals with autism were tested for Fragile-X syndrome using the AmplideX^TM^ *FMR1* PCR kit from Theradiag. There were no individuals with “pre-mutation” or “full-mutation” of CGG repeats in the 5’ UTR region of the fragile X mental retardation-1 (*FMR1*) gene.

### Copy-number variants (CNVs)

CNVs were identified from both SNP genotyping and WES data. Quality controls were the following: call rate > 0.99, standard deviation of the Log R ratio < 0.35, standard deviation of the B allele frequency < 0.08 and absolute value of the wave factor < 0.05. CNVs were detected by both PennCNV(70) and QuantiSNP(71) algorithm using the following filters: >= 3 consecutive probes, CNV size > 1kb and CNV detection confidence score >= 15. CNV detections from PennCNV and QuantiSNP were merged using CNVision(17). CNVs with CNVision confidence score < 30, CNV size < 50 kb, overlap > 50% with segmental duplication or known large assembly gaps (greater than 150 kb) or copy number = 2 in pseudo autosomal regions (PARS) in males were filtered out. CNV annotations were performed using ANNOVAR (72) and CNV frequencies in Faroese and in database of genomic variant cohorts (DGV, http://dgv.tcag.ca/dgv/app/home) were assessed using in house python scripts based on reciprocal overlap >= 80%. We also detected CNVs from the WES sequencing data using the XHMM software(73). CNVs with QSOME score < 90, number of targets < 5, or overlap > 50% with segmental duplication or known large assembly gaps (greater than 150 kb) were filtered out. CNV annotations were performed using ANNOVAR (72) and CNV frequencies in Faroese were assessed using in house python scripts based on reciprocal overlap ≥ 80%. *De novo* and inherited CNVs were validated by visual inspection using SnipPeep (http://snippeep.sourceforge.net/).

### Gene-set lists and prioritization of variants

Three gene-set lists were used : (i) “SFARI genes” (n=990) that includes genes implicated in autism (15) (Simons Foundation Autism Research Initiative gene database - https://gene.sfari.org/); (ii) “pLI > 0.9 genes” that includes genes with strong probability of being loss-of function intolerant (n=3,230) (74); (iii) “Brain genes” that includes genes specifically or strongly expressed (above 1 Standard Deviation) in fetal or adult human brain using data from Su et al (n=3,591)(75).

A combination of approaches was used to prioritize the genes and to estimate the deleterious effect of a variant. We prioritized genes using gene sets (SFARI genes, pLI >= 0.9 and Brain genes, previously defined). We prioritized Likely Gene Disruptive (LGD) variants (stopgains, splice site variants, frameshift indels) over missense variants or synonymous variants. Additionally, we used the CADD score (14) (a CADD >= 30 means that the variants belong to the 0.1% most deleterious variants) to assess the deleterious effect missense variants. Minor allele frequency (MAF) was estimated in the general population from the gnomAD database(74). In order to filter out common variants that was not listed in gnomAD, we also excluded variants that were present in more than 15% of our Faroese control cohort. For the detection of deleterious homozygous variants, we kept only LGD and MIS30 with MAF <1%.

### Burden analysis

Rare variant association studies (MAF<5%) were performed using EPACTS v3.2.6 (https://genome.sph.umich.edu/wiki/EPACTS). Prior association analysis, variants identified by WES were filtered using VCFtools (http://vcftools.sourceforge.net/man_latest.html) with the following metrics: minimum genotyping quality ≥ 30, min depth of coverage ≥ 10, maximum of missing data ≤ 10, no InDel (small insertion or deletion), only bi-allelic sites and no site failing Hardy Weinberg equilibrium test (*p* < 10^−6^). The annotation of the variants was done using EPACTS and the variants included in the Gene-wise association analyses were non-synonymous, essential splice site, normal splice site, start loss, stop loss and stop gain variants. Logistic Score Test (“b.score” in S1 Table) was used to test single variant association (n_Cases_=36; n_Controls_=107; n_Variants_=155,284). For Gene-wise tests, we used two approaches (including n_Cases_=36; n_Controls_=107 and n_groups_= 15,005): (i) collapsing burden test using EMMAX (Efficient Mixed Model Association eXpedited (76), “CMC-EMMAX” in S1 Table) and (ii) Optimal SNP-set sequence Kernel Association Test (“SKAT-O” in S1 Table). The advantage of the CMC-EMMAX is that this test is accounting for population structure and high relatedness between individual (based on kinship matrix). The advantage of SKAT is that this test is particularly powerful in the presence of protective and deleterious variants and null variants. For both Gene-wise tests, a 10^−6^ ≤ MAF ≤ 0.05 was used.

## Acknowledgements

We would like to thank all the participants of the epidemiological and genetic cohorts, the “progeny” database, Julien Fumey for his R tutorial (https://bioinfo-fr.net/creer-sa-carte-geographique-avec-r) and Fabrice de Chaumont for his help with python.

## Funding

This work was supported by the Institut Pasteur; Centre National de la Recherche Scientifique; the Assistance Publique – Hôpitaux de Paris; the University Paris Diderot; the Simons Foundation; the Fondation pour la Recherche Médicale [DBI20141231310]; the European Commission Horizon 2020 [COSYN]; The human brain project; the European Commission Innovative Medicines Initiative [EU-AIMS no. 115300]; the Cognacq-Jay foundation; the Bettencourt-Schueller foundation; the Orange foundation; the FondaMental foundation; the Conny-Maeva foundation; and the Agence Nationale de la Recherche (ANR) [SynPathy]. This research was supported by the Laboratory of Excellence GENMED (Medical Genomics) grant no. ANR-10-LABX-0013, Bio-Psy and by the INCEPTION program ANR-16-CONV-0005, all managed by the ANR part of the Investment for the Future program. The funders had no role in study design, data collection and analysis, decision to publish, or preparation of the manuscript.

